# Intranasal oxytocin reduces pre-courtship aggression and increases paternal response in California mice (*Peromyscus californicus*)

**DOI:** 10.1101/2021.06.20.449160

**Authors:** Caleigh D. Guoynes, Catherine A. Marler

## Abstract

Oxytocin (OXT) is a neuropeptide that can facilitate prosocial behavior and decrease social stress and anxiety. We investigated whether acute pulses of intranasal (IN) OXT influenced social behavior during social challenges that are likely to occur throughout the lifespan of a wild mouse. To test this, we examined the acute effects of IN OXT in the male California mouse (*Peromyscus californicus*), a monogamous, biparental, and territorial rodent, using a within-subjects longitudinal design. Social challenges included a pre-courtship male-female encounter conducted during the initial aggressive and not the following affiliative phase of courtship, same-sex resident intruder test, and parental care test, with each test and dose separated by at least two weeks. Males were treated with intranasal infusions of 0.8 IU/kg OXT or saline controls 5-min before each behavioral test, receiving a total of three treatments of either IN OXT or saline control. We predicted that IN OXT would 1) decrease aggression and increase affiliation during the pre-courtship aggression phase, 2) increase aggression during resident intruder paradigms and 3) increase paternal care and vocalizations during a paternal care test. As predicted, during pre-courtship aggression with a novel female, IN OXT males displayed less contact aggression than control males, although with no change in affiliative behavior. However, post-pairing, during the resident intruder test, IN OXT males did not differ from control males in contact aggression. During the paternal care test, IN OXT males were quicker to approach their pups than control males but did not differ in vocalizations produced, unlike our previous research demonstrating an effect on vocalizations in females. In summary, during pre-courtship aggression and the paternal care test, IN OXT promoted prosocial approach; however, during the resident intruder test IN OXT did not alter social approach. These data suggest that IN OXT promotes prosocial approach specifically in social contexts that can lead to affiliation.

**HIGHLIGHTS:**

- IN OXT attenuates male aggression during pre-courtship encounters
- IN OXT does not attenuate male aggression during resident intruder encounters
- IN OXT increases paternal responsiveness during a paternal care challenge
- IN OXT in fathers does not influence total paternal care or vocalizations

**GRAPHICAL ABSTRACT:** 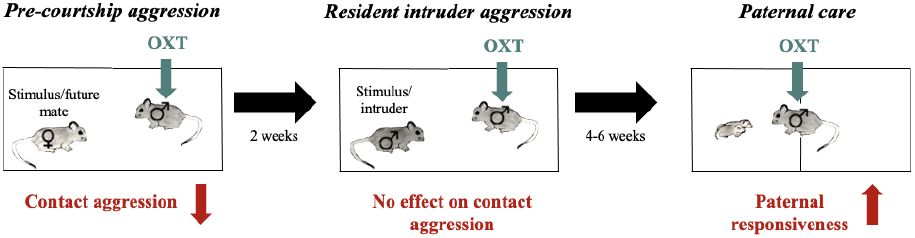

## 1. Introduction

In social species, interactions can be altered based on their life history stage and environment. Throughout the lifespan, social species encounter many different types of social interactions and must respond appropriately to these social interactions to acquire and maintain resources, mating opportunities, and reproductive fitness. One significant question is determining the mechanisms underlying how animals alter their social responses based on social and environmental context and life stage. Endogenous hormone and neuropeptide levels are important for biobehavioral feedback and to help animals respond appropriately to various social interactions. Oxytocin (OXT), a neuropeptide hormone, is a neuromodulator that may be important for weighing social salience and determining appropriate behavioral response to social stimuli (Shamay-Tsoory & Abu-Akel, 2016; Parr et al., 2018; Yao et al., 2018; Johnson et al., 2017; Egito et al., 2020). Previous studies on OXT show its significant effects on prosocial affiliative behaviors such as trust, social bonding, social recognition, and anxiolytic behavior in both human and animal models (Theodoridou et al., 2009; Kosfeld et al., 2005; Ring et al., 2006; Bales et al., 2003; Blocker et al., 2015; Guestella et al., 2008). In addition to increasing affiliative behaviors, OXT is involved in aggressive behaviors. In humans, OXT can increase envy, schadefreude, defensive but not offensive aggression toward a competing out-group, and domestic violence in men prone to aggression (Shamay-Tsoory et al. 2009; Bethlehem et al., 2015; De Dreu et al., 2016; De Dreu et al., 2010; DeWall et al. 2014). OXT is also associated with increased mate guarding in rats (Holley et al., 2015), prairie voles (Bales & Carter 2003), and marmoset monkeys (Cavanaugh et al., 2018). Furthermore, OXT is associated with increased maternal aggression toward potential predators (Bosch & Neumann 2012). In canines, OXT also increases aggression towards owners but not strangers during a threatening approach test (Hernadi et al., 2015). These data on the role of OXT on affiliative and aggressive behavior support the hypothesis that social salience and social context are important cues influencing the behavioral effects of OXT. Based on these studies, OXT would be expected to decrease aggression and increase affiliative behavior when a male-female pair is introduced and increase aggression by a resident towards an intruder.

Throughout an animal’s lifetime, OXT levels change in response to certain life events such as early life experience, pair bonding, intrasexual aggression, and parenting. This is especially true for monogamous and parental species that require flexibility in response to group membership. In prairie voles, the function of OXT can be altered in response to previous social neglect by their mother during the neonatal period (Bosch and Young, 2017). Prior to mating, OXT increases affiliative contact with familiar females (Cho et al. 1999; Bales et al., 2013) and increases speed of pair bonding in females (Williams et al., 1994; Young & Wang, 2004). Postmating, OXT enhances aggression in prairie voles during encounters with same-sex conspecifics (Winslow et al. 1993). In California mice, OXT plasma levels increase in expectant fathers, decrease in fathers, and are disrupted when the male is separated from his mate and pups (Gubernick et al., 1995). These rodent studies in prairie voles and California mice suggest that social experience may drive important changes to the OXT system. These studies further enhance expectations for OXT to increase paternal behavior.

To mimic the natural pulses of OXT that may occur during these different social contexts and challenges, acute intranasal OXT (IN OXT) can be used. Previous studies in rodents have shown that IN OXT alters behavior within 5-min of administration (Bales et al., 2013) and can have behavioral effects that persist for 30-50 min after administration (Carter & Wilkinson, 2015). Daily chronic doses of IN OXT induce long-term modifications to the OXT system (Bales et al., 2013; Guoynes et al., 2018; Del Razo et al., 2020); however, single doses spread out across weeks are presumably less likely to have carry-over effects across tests (Huang et al., 2014).

The California mouse (*Peromyscus californicus*) is a strictly monogamous, biparental rodent species well-suited to examine how OXT modulates vocal production and social behavior across different life stages. California mice show aggression toward unfamiliar conspecifics (e.g. Rieger et al. 2018) including opposite-sex conspecifics (e.g. e.g.Pultorak et al., 2018). During pre-courtship aggression with an unfamiliar conspecific, there is a period of assessment and often aggression (Gleason & Marler, 2010) that we will refer to as the pre-courtship aggression phase. Most of this aggression is in the form of non-contact aggression such as chasing and lunging, but the aggression can escalate to contact forms of aggression such as wrestling. Based on previous experience pairing female and male California mice in the lab, most prospective pairs show some form of aggression (i.e. lunging, chasing) but fewer pairs show contact aggression (i.e. wrestling) (Gleason & Marler, 2010). Once paired, female and male California mice form strong, reliable pair bonds but will still show reliable aggression toward unfamiliar conspecifics (Bester-Meredith & Marler, 2001; Trainor & Marler, 2001; Bester-Meredith & Marler, 2007); such aggression is decreased by an antagonist (V1a) to vasopressin (Bester-Meredith et al. 2005), a similar neuropeptide that is often positively associated with aggression. The period of pre-courtship aggression in the California mice is significantly longer than in other monogamous animal models such as the prairie vole. While prairie voles mate within the first 41 hrs of being paired (Witt et al., 1988), California mice mate 7-14 days after being paired (Bester-Meredith et al., 2003; Trainor et al., 2001; Gleason & Marler 2010). This longer period of courtship may reflect a longer assessment period for potential mates, as expected in a monogamous species. The first litter of pups is typically born between six and eight weeks after the initial pre-courtship aggression. Once pups are born, both fathers and mothers engage in parental care (Bester-Meredith & Marler, 2001; Bester-Meredith & Marler, 2003; Lee & Brown 2002; Trainor et al., 2003; Trainor & Marler, 2003; Marler et al., 2003; Lee et al., 2007; Frazier et al., 2006; Becker et al., 2010; Gleason & Marler, 2010; Bester-Meredith & Marler, 2012; Johnson et al., 2015; Rieger et al., 2019; Guoynes & Marler, 2021).

California mice also have a diverse, well-characterized repertoire of ultrasonic vocalizations (USVs) including simple sweeps, complex sweeps, syllable vocalizations, barks, and pup whines (Briggs et al. 2011; Kalcounis-Rueppell et al., 2006; Pultorak et al., 2015; Rieger & Marler, 2018; Guoynes & Marler, 2021). A previous study in mother-offspring interactions demonstrated that the primary call types observed were maternal simple sweeps and pup whines; maternal simple sweeps correlated with both maternal care and pup whines (Guoynes & Marler, 2021). Similar to the prevalence of call types in mother-offspring interactions, preliminary recordings between fathers and pups indicated that the primary call types from fathers and pups were also paternal simple sweeps and pup whines, respectively. Moreover, OXT stimulated production of maternal sweeps (Guoynes & Marler 2021). Based on this, we predicted a similar response to OXT in fathers involving simple sweeps and pup whines. It is important to note that paternal simple sweeps and pup whines have also been recorded in other social contexts (Guoynes & Marler, 2021; Rieger et al., 2019; Pultorak et al., 2015; Pultorak et al., 2017). Because we were not manipulating the OXT system in the pups, we did not expect to see an effect of OXT on pup whine USVs.

In the current study, we aimed to address whether acute pulses of IN OXT alter an animal’s response to social challenges. We hypothesized that 1) during the pre-courtship aggression phase, IN OXT would reduce aggression, specifically the escalation to contact aggression (i.e., wrestling) in male-female aggression and increase affiliative behavior, 2) during resident intruder paradigms IN OXT would increase aggression towards an intruding male and 3) during a parental care test, similar to the effects in mothers, IN OXT would have a positive effect on paternal care and paternal vocalizations.

## 2. Methods and Materials

### 2.1. Animals

University of Wisconsin-Madison Institutional Animal Care and Use Committee approved this research. We used 24 male *P. californicus* aged 5–10 months. They were group-housed (2–3 per cage; 48 × 27 × 16 cm) under a 14L: 10D light cycle with lights off at 4:00pm. Animals were maintained in accordance with the National Institute of Health Guide for the Care and Use of Laboratory Animals. Males were randomly assigned to either the saline control group (N=12) or the OXT group (N=12). The OXT group received three total doses of OXT and the saline group received three total doses of saline (one dose given 5-min before each behavioral test) over eight weeks. For pair bond initiation, 24 female mates unrelated by at least two generations were randomly assigned to the focal test males. For the resident intruder test, 24 unrelated male intruders were randomly assigned to the focal test males. During the paternal care test, pup number across treatments was very similar such that the average number of pups for fathers in the saline control condition was 2.13 ± 0.23 (mean ± SE), and average number of pups for fathers in the OXT condition was 2.25 ± 0.16 (**S. Table 3**).

### 2.2. Intranasal Oxytocin Preparation

Male mice were infused intranasally with either sterile saline or IN OXT (0.8 IU/kg) (Bachem, Torrance, California) (Guoynes & Marler, 2021). The IN OXT dose is equivalent to doses used in other animal models (Bales et al. 2014; Guoynes et al. 2018; Murgatroyd et al. 2016) and similar to weight-adjusted doses used in clinical studies examining the effects of IN OXT on social deficits in autism (Bales et al., 2013). IN OXT was dissolved in saline and prepared in one batch that was aliquoted into small plastic tubes and frozen at 20°C. IN OXT was defrosted just prior to administration. A blunt cannula needle (33-gauge, 2.8 mm length; Plastics One, Roanoke, Virginia) was attached to cannula tubing, flushed, and filled with the compound, then attached to an airtight Hamilton syringe (Bachem, Torrance, California). The animal was scruffed and 25 uL of compound was expelled dropwise through the cannula needle and allowed to absorb into the nasal mucosa (~10-20 seconds). One person conducted all IN OXT administrations throughout the entire procedure to maintain consistency in handling and IN OXT infusion. We chose to use the method of intranasal administration of IN OXT for two primary reasons. (1) IN OXT is used in clinical studies and is less invasive, does not require special transporters for the molecule, and is presumed to be less stressful compared to intracerebroventricular (Talegaonkar & Mishra 2004). (2) IN OXT shows similar behavioral effects as centrally administered OXT, increases CSF and plasma concentrations of OXT, and reaches the relevant brain areas in both humans and animal models (Neumann et al., 2013; Striepens et al., 2013; Lee et al. 2018; Oppong-Damoah et al., 2019; Lee et al., 2020). Several studies have also shown changes in plasma OXT concentrations that peak between 15 to 30-min post-administration (Freeman et al., 2016; Gossen et al., 2012). These results suggest IN OXT passes through the blood-brain barrier to exert central effects. In California mice, behavioral effects of IN OXT are consistent with the outcomes of central OXT manipulations suggesting that IN OXT is reaching the brain (Duque-Wilckens et al. 2018, 2020). Other studies indicate that some of the effects of IN OXT are acting through peripheral mechanisms (Churchland & Winkielman, 2012; Quintana et al., 2015; Leng & Ludwig, 2016). Regardless of whether IN OXT is directly targeting the brain, is acting through peripheral mechanisms, or a combination of both, IN OXT has been shown to rapidly alter social behavior in adult California mice (Steinman et al., 2016).

### 2.3. Behavioral Tests

Throughout the experiment, all researchers administering treatments and handling animals were blind to treatment condition. For each test, the same researcher administered all intranasal treatments to reduce variance across handling and administration.

#### Pre-courtship aggression test

Male California mice aged 5-10 months were removed from their home cage (48 × 27 × 16 cm) and given 25 uL of 0.8 IU/kg OXT or saline. Immediately after treatment, each male was placed in a new home cage (48 × 27 × 16 cm) with fresh bedding. 5-min after the dose of OXT or saline, a novel, unrelated female was placed into the new home cage. Their interaction was videotaped for 10-min (**Fig. 1A**). After the recording, the male and female continued to be housed together for the remainder of the experiments.

**Figure 1.**
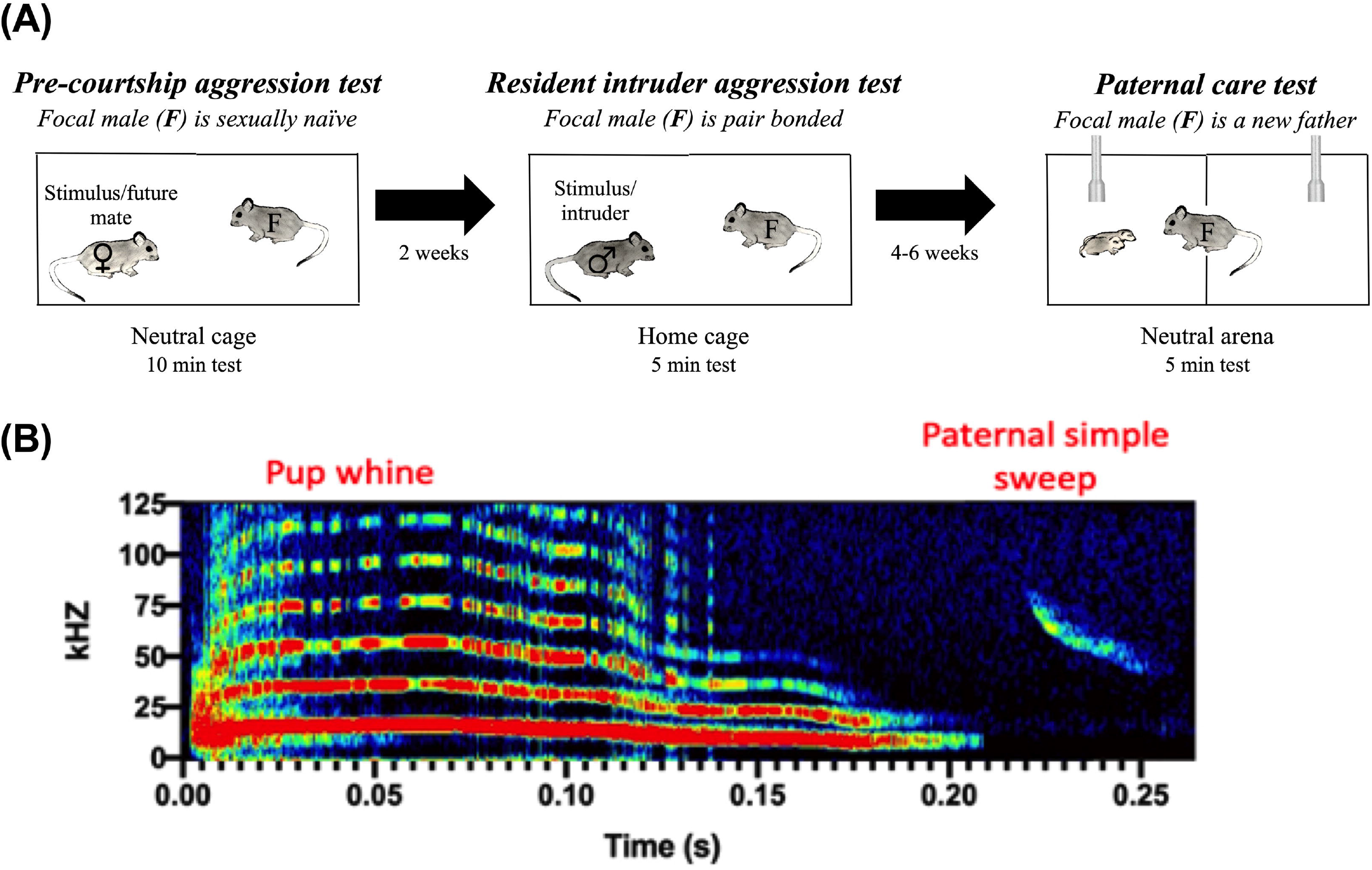
Experimental design. **(A)** Timeline of the three behavioral tests throughout the longitudinal study. **(B)** Representative pup whine and paternal simple sweep USVs. Pup whines have multiple harmonics, a peak frequency around 20 kHz, and downward modulation at the end of the call that distinguish these calls from adult syllable vocalizations. Paternal simple sweeps have short downward-sweeping vocalizations that sweep through multiple frequencies, typically between 80 kHz and 40 kHz.

#### Resident intruder test

We continued to use the same male and female pairs as in the pre-courtship aggression test above, but 14 days after being paired. Residency in the home cage was established by housing the mice in the same home cage for 6 consecutive days. This is more than sufficient time to establish residency in males (Bester-Meredith et al., 1999; Marler et al., 2003; Fuxjager et al., 2010; Zhao et. al 2014). Immediately before testing, female pair mates were removed from the home cage and placed in a new home cage with fresh bedding adjacent to the old home cage with soiled bedding (each 48 × 27 × 16 cm). Male pair mates were given 25 uL of 0.8 IU/kg OXT or saline (same treatment as they received in the pre-courtship aggression test) and placed back in their home cage with soiled bedding. 5-min after administration of OXT, an unrelated, novel male was placed on the far side of the resident’s cage. Their interaction was recorded for 5-min (**Fig. 1A**). After the test, the novel male was removed and placed back in his home cage and then the resident male given OXT or saline was removed and placed into the clean home cage with his female pair mate.

#### Paternal care test with ultrasonic vocalizations (USVs)

This test used the same male and female pairs as in the pre-courtship aggression test and resident intruder test (above) and was conducted three to six weeks after the resident-intruder test—on the first or second day after their first litter was born. Pairs were monitored and checked for pups daily. Testing occurred within 48 hrs of the pups being born during a stage of postpartum estrous. The pups were removed from the mother, and the mother was placed in a new home cage with some soiled bedding from the home cage. Next, the father and pups in their home cage were transferred from the mouse housing room to a behavior testing room capable of recording USVs. This procedure is similar to paradigms previously used in the lab (Guoynes & Marler, 2021; Pultorak et al., 2015; Rieger & Marler, 2018). Testing was done in a custom arena split into two equally sized chambers (45.0 cm × 30.0 cm × 30.0 cm) and contained two symmetrically located circular openings (3.8 cm in diameter, center of opening 7 cm from the side wall) covered by a wire mesh. Ultrasonic microphones (described below) were placed on each side of the divider. One side of the divider was designated to the focal male, the other to the pup(s). This setup allowed visual, auditory, and olfactory communication between pups and their father, but restricted physical contact between individuals until the mesh wire was removed. In the testing room, fathers were given a third dose of either 25 uL of 0.8 IU/kg OXT or saline (same treatment as they received in the pre-courtship aggression test and aggression test) and placed back into their home cage for 5-min (**Fig. 1A**). At the end of the 5-min waiting period, the pups were moved into the side of the testing chamber near the door, and the fathers were moved into the chamber closest to the wall. They interacted through the mesh divider intact for the first 3-min, then the divider was removed, and the fathers and pups could physically interact for an additional 5-min. Vocalizations and video were recorded for the entire 8-min period. These time periods were chosen because they minimized the time that the pups were away from their mother but allowed enough time to quantify behavioral differences in retrievals.

### 2.4. Behavior Quantification

All behavior videos were scored twice: once each by two independent observers blind to treatment and in a random order. Scores between observers had to be at least 85% similar and scores between the two observers were averaged for the final output used in statistical analysis. For an ethogram describing these different behaviors

### 2.5. Ultrasonic Vocalization Analysis

Techniques used for recording were similar to those previously used in our laboratory (Pultorak et al. 2017; Rieger & Marler 2018; Guoynes & Marler 2021). USVs were collected using two Emkay/Knowles FG series microphones capable of detecting broadband sound (10–120 kHz). Microphones were placed at the far ends of each of the two chambers. Microphone channels were calibrated to equal gain (−60 dB noise floor). We used RECORDER software (Avisoft Bioacoustics) to produce triggered WAV file recordings (each with a duration of 0.5 s) upon the onset of a sound event that surpassed a set threshold of 5% energy change (Kalcounis-Rueppell et al., 2010). Recordings were collected at a 250 kHz sampling rate with a 16-bit resolution. Spectrograms were produced with a 512 FFT (Fast Fourier Transform) using Avisoft-SASLab Pro sound analysis software (Avisoft Bioacoustics). The only USVs found in these recordings were pup whines and paternal simple sweeps. Pup whines have a peak frequency around 20 kHz (Johnson et al., 2017; Kalcounis-Rueppell et al., 2018a) and the typical downward modulation at the end of the call often distinguishes these calls from adult syllable vocalizations (Guoynes & Marler, 2021; Nathaniel Rieger, Jose Hernandez, & Catherine Marler, unpublished) (**Figure 1B**). The lower frequencies in the pup whine can also be heard by human ears (below the ultrasonic range). Paternal simple sweeps were categorized by short downward-sweeping vocalizations that sweep through multiple frequencies, typically between 80 kHz and 40 kHz (Kalcounis-Rueppell et al., 2018b) (**Figure 1B**). It is extremely rare for pups to produce simple sweep USVs during PND 0-4 (Rieger, N. S., Hernandez, J. B., and Marler, C. M., unpublished). When young pups produce simple sweeps, they are produced much faster and present completely vertical on the spectrogram (Johnson et al., 2017). This makes these rare pup simple sweeps easy to distinguish from the slower adult simple sweep USVs (**Fig. 1B**). Because of their different spectrogram and acoustic properties, all USVs could be categorized and counted by combined visual and auditory inspections of the WAV files (sampling rate reduced to 11,025 kHz, corresponding to 4% of real-time playback speed).

### 2.6. Data Analysis

For each behavioral test, nonparametric Mann-Whitney tests were conducted to compare the outcomes between saline control and OXT males. In the pre-courtship aggression test, one OXT mouse was dropped from the analysis because he escaped from the apparatus just prior to testing. Final group size analyzed for the pre-courtship aggression test was N=12 for control males and N=11 for OXT males. In the resident intruder test, final group size analyzed for the pre-courtship aggression test was N=12 for controls and N=12 for OXT males. In the paternal care test, three pairs were removed from behavioral analyses due to accidental deleting of the behavior videos (1 control male, 2 OXT males), and 5 were not tested because of either infanticide or not producing pups within eight weeks of pairing. Final group size analyzed for the behavioral and USV components of the paternal care test was N= 8 for controls and N=8 for OXT.

Correlations between paternal care and USVs were conducted using the program R. To assess for mediation by IN OXT in the relationships between (a) paternal USVs and paternal behavior and (b) paternal behavior and pup USVs, a multivariate comparison was used. Factors included in the model were treatment condition and the interaction between treatment and paternal behavior`(e.g. [Paternal behavior] ~ [Paternal USV] + [treatment])`.

Significance level was set at p < 0.05 for all analyses and all tests were two-tailed. All reported p-values were corrected using Benjamini-Hochberg false discovery rate corrections to control for multiple comparisons when effect of an X variable was tested for a relationship with multiple Y variables. False discovery rate was set at five percent.

## 3.0 Results

### 3.1. Pre-courtship aggression test

To determine whether IN OXT influenced escalation to contact aggression during pre-courtship aggression, we assessed number of wrestling bouts in male mice given IN OXT versus saline. We found that OXT decreased the proportion of wrestling bouts out of all aggressive behaviors between the male and female during the first 10-min of pre-courtship aggression (U=33, z-*score*=2.00, *p*<0.05) (**Fig. 2A**). Lunging aggression levels made up a relatively small proportion of the aggressive behaviors in both control and OXT males; however, differences arose in proportion of wrestling aggression (highest in control males) and chasing aggression (highest in OXT males) (**Fig. 2B**). Levels of non-contact aggression were relatively similar across groups (lunging aggression: CTRL=1.29± 1.36 and OXT=0.45± 0.37; chasing aggression: CTRL=10.76± 3.73 and OXT=12.80± 3.82) (**S. Table 1**). The biggest difference between treatment groups was amount of time spent engaged in contact aggression (wrestling aggression: CTRL=11.58± 6.22 and OXT=0.77± 0.50) (**S. Table 1**). Thus, the difference in proportion of wrestling of aggression between CTRL and OXT is being driven by time spent wrestling vs. time spent chasing. Other behaviors we did not predict would be affected by IN OXT such as social investigation (body and anogenital sniffing) and activity (autogrooming, rearing) were measured but not statistically analyzed (**S. Table 1**).

**Figure 2.**
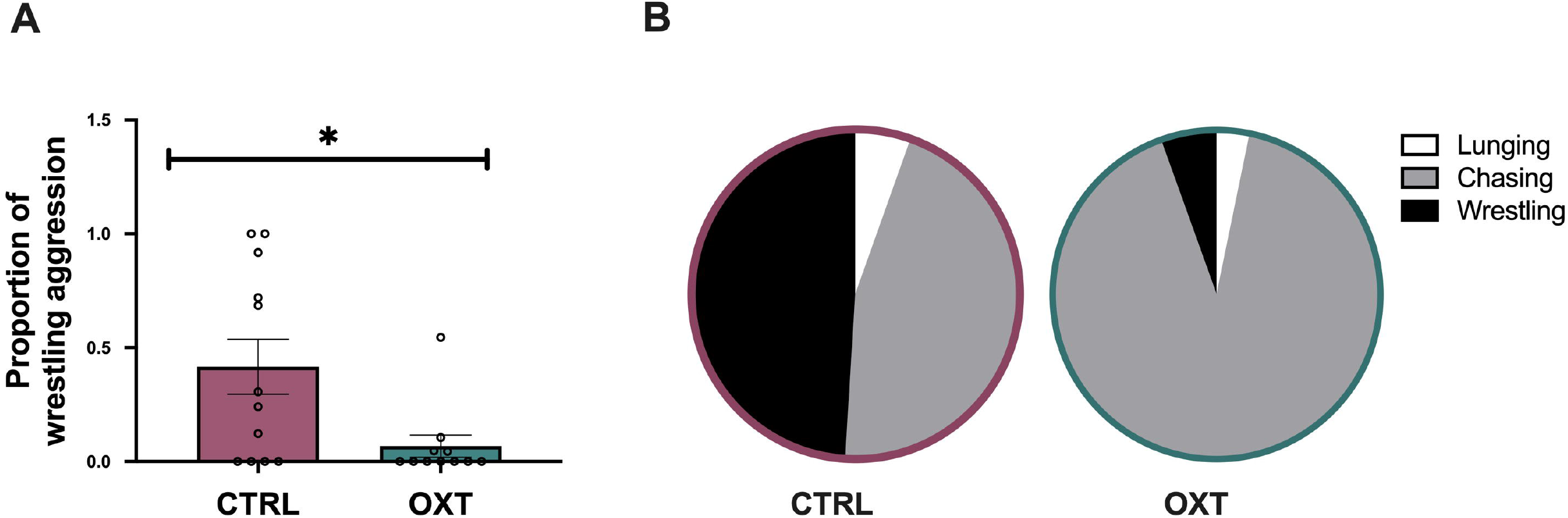
Pre-courtship aggression test. Males given OXT had a significantly smaller proportion of wrestling than control males during the first 10 min of courtship. **(B)** Pie chart showing escalating aggressive behavior (from light: low escalation, to dark: high escalation). *p<0.05 for differences between control and OXT.

### 3.2. Resident intruder aggression test

To determine whether IN OXT influenced escalation to contact aggression during a resident intruder test, we assessed the number of wrestling bouts in males given IN OXT versus saline. Unlike the pre-courtship aggression test, we found that IN OXT did not significantly influence number of wrestling bouts between the males during a 5-min resident intruder test (U=63.50, z-*score*=0.46, *p*=0.637) (**Fig. 3A**). Similar to the pre-courtship aggression test, lunging aggression levels made up a relatively small proportion of the aggressive behaviors in both control and OXT males (**Fig. 3B**). Both chasing and wrestling aggression made up approximately equal proportions of aggressive behavior in the resident intruder aggression test (**Fig. 3B**). Levels of all types of aggression were relatively similar across groups (lunging aggression: CTRL=2.25± 1.00 and OXT=1.63± 0.71; chasing aggression: CTRL=13.63± 5.52 and OXT=11.28± 5.86; wrestling aggression: CTRL=11.47± 4.52 and OXT=10.69± 4.63) (**S. Table 2**). Other behaviors we did not predict would be affected by IN OXT such as social investigation (body and anogenital sniffing) and activity (autogrooming, rearing) were measured but not statistically analyzed (**S. Table 2**).

**Figure 3.**
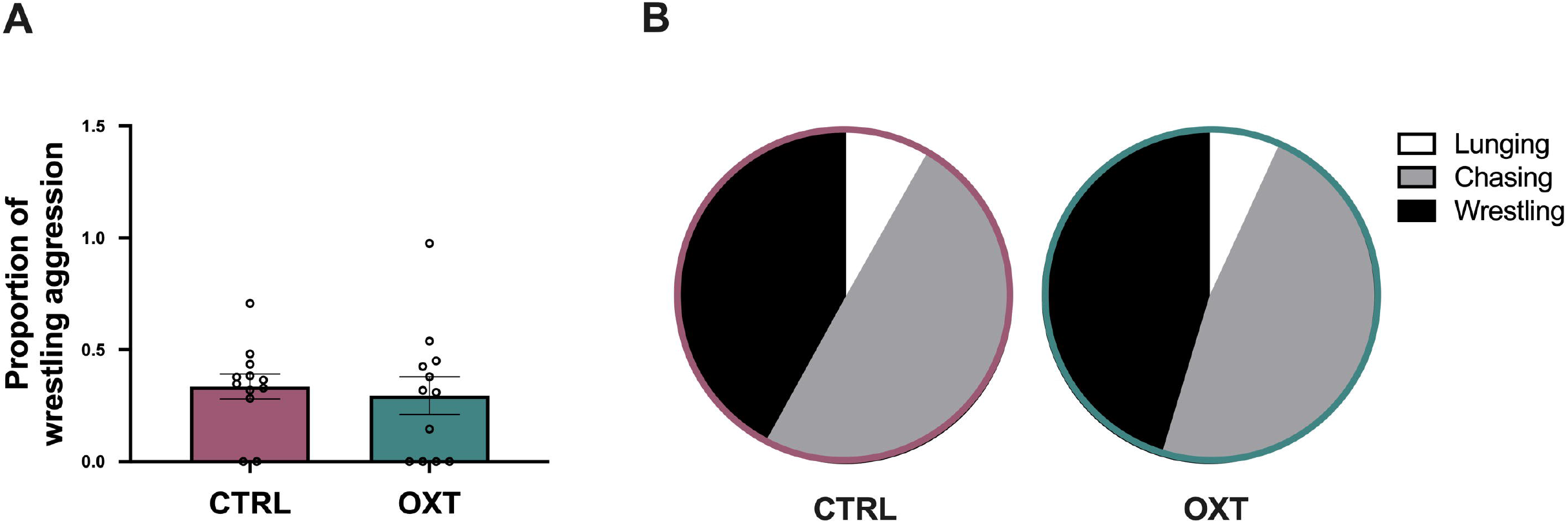
Resident intruder aggression test. **(A)** OXT and control males showed no difference in proportion of wrestling during a 5-min resident intruder encounter. **(B)** Pie chart showing escalating aggressive behavior (from light to dark). *p<0.05 for differences between control and OXT.

### 3.3. Paternal care test with ultrasonic vocalizations (USVs)

To determine whether IN OXT would influence behavior during a paternal care challenge we assessed latency to approach pups, pup huddling, and paternal simple sweep USVs in fathers given IN OXT versus saline. Fathers given IN OXT were significantly faster at approaching their pups after a brief separation (*U*=10.50, z-*score*=2.21, *p*<0.05) (**Fig. 4A**). Despite initial differences in paternal care response, there were no differences between IN OXT and control males in total time huddling (*U*=22.50, z-*score*=-0.95, *p*=0.34) (**Fig. 4B**) or licking pups (*U*=20, z-*score*=-1.21, *p*=0.22) (**Fig. 4C**). There was one father in the control group that showed much more paternal care than other control fathers, however, this father was not a Grubb’s outlier for paternal care measures. Even if this father is removed from the analysis, the difference between control and OXT is not significant for huddling (*U*=14.50, z-*score*=1.50, *p*=0.13) (**Fig. 4B**) or licking (*U*=12, z-*score*=1.79, *p*=0.07) (**Fig. 4C**). Neither IN OXT or control fathers engaged in any retrieval behavior throughout the test, so this type of paternal care was not analyzed (**S. Table 3**). There were no differences in number of pups across treatments groups (CTRL=2.13± 1.84 1.84; OXT=2.25± 1.54). Other behaviors related to activity (autogrooming, freezing, rearing) were measured but were not included in the statistical analyses because we did not have *a priori* predictions for these behaviors during the paternal care test (**S. Table 3**).

**Figure 4.**
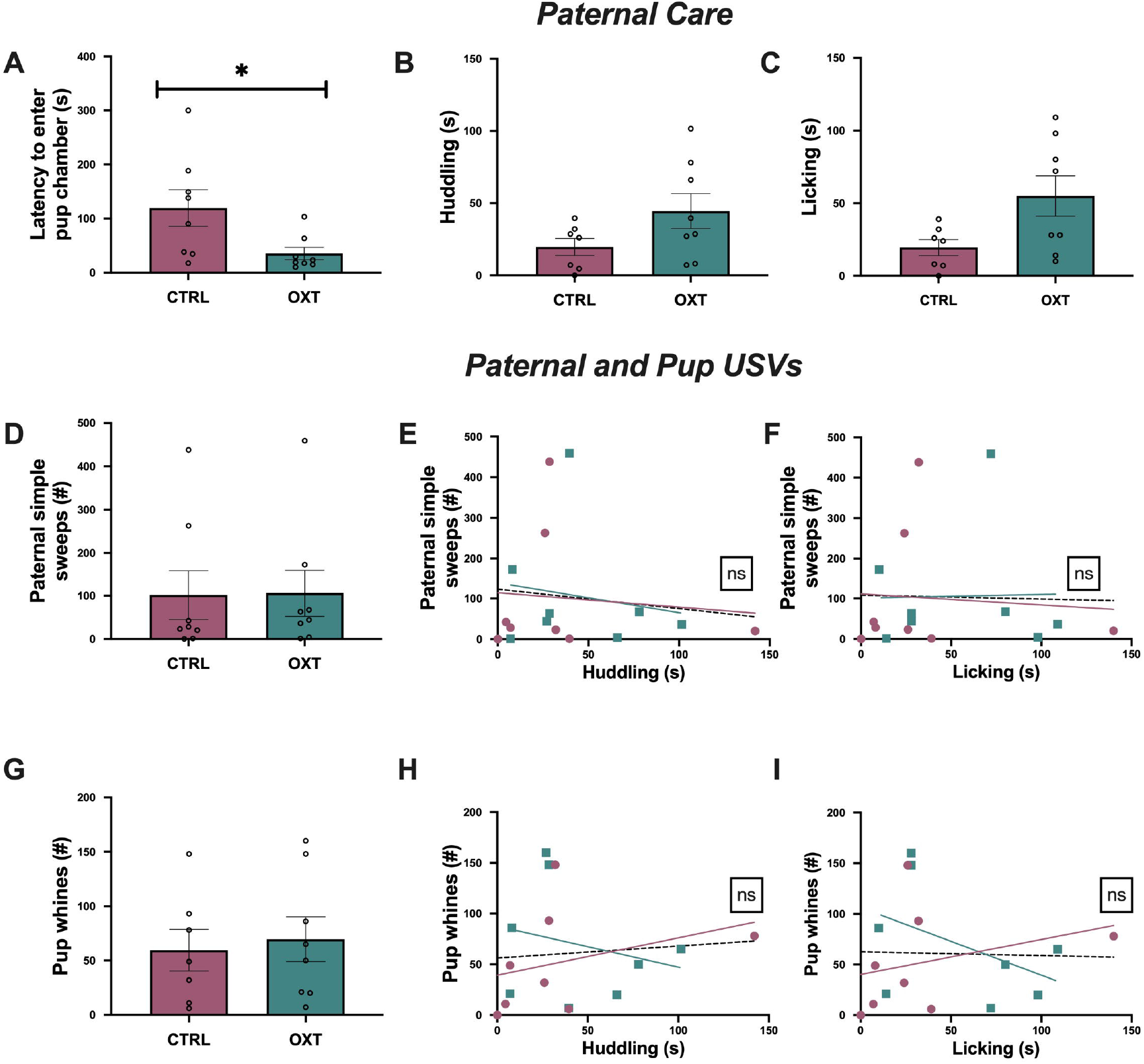
Paternal care test. OXT males had shorter latencies to approach their pups than control males **(A)**. OXT males did not show significant differences in huddling **(B)** or licking **(C)** behavior. **(D)** Males given OXT did not make more simple sweeps than control males. Paternal simple sweeps did not correlate with **(E)** huddling or **(F)** licking. **(G)** Pups with OXT versus control fathers showed no differences in number of pup whines produced. There were no correlations between pup whines and **(H)** huddling or **(I)** or licking. *p<0.05 for differences between control and OXT.

Next, we assessed whether IN OXT would influence paternal and/or pup USVs behavior during a paternal care challenge. We assessed number of paternal simple sweeps and number of pup whines produced and their correlations with the two types of paternal care observed, huddling and licking. Fathers given IN OXT did not produce more simple sweeps than controls (*U*=23.50, z-*score*=-0.84, *p*=0.40) (**Fig. 4D**). There were also no differences in number of pup whines produced in offspring of IN OXT versus control fathers (*U*=24.50, z-*score*=0.35, *p*=0.72) (**Fig. 4E**).

Lastly, we examined the relationship between paternal care and paternal and pup USVs and any interactions with OXT treatment. Using a multivariate model controlling for the effects of treatment, we found no main effects of paternal simple sweeps on huddling (F_2,16_=0.21, *p*=0.65, η^2^=0.016) (**Fig. 4H**) or licking (F_2,16_=0.01, *p*=0.91, η^2^=0.00) (**Fig. 4J**). Similarly, we found no main effects of pup whines on huddling (F_2,16_=0.05, *p*=0.81, η^2^=0.00) (**Fig. 4I**) or licking (F_2,16_=0.07, *p*=0.80, η^2^=0.00) (**Fig. 4K**).

## 4. Discussion

Our study assessed the response of male California mice to different challenges that would naturally occur during their lifespan. During contexts in which the social stimuli had the potential to become part of the in-group, a male-female bonded pair, OXT administered to the male promoted prosocial approach through reduced aggression. In contrast, during the resident-intruder aggression test, the social stimuli did not have the potential to become part of the ingroup in a strongly territorial species, and OXT did not promote prosocial approach. Finally, in the paternal behavior test, OXT increased paternal motivation to approach pups in this biparental species. We speculate that OXT may function to promote social approach only in contexts that are or are likely to be affiliative-prone.

In the monogamous and territorial California mice, when virgins encounter an unfamiliar individual of the opposite sex, there is both an aggressive response to an unfamiliar conspecific, and possibly novelty, and a potential for pair bond formation. During the initial 10-min of this interaction, only aggressive behavior was exhibited, with no signs of affiliative behavior characteristic of later stages of courtship (Gleason & Marler, 2010) or as they are bonding (Pultorak et al., 2017); also similar to the behavioral sequence seen in research with other species between male and females prairie voles (Williams et al., 1992; Carter et al., 1995; Cho et al., 1999; Willett et al., 2018; Harbert et al., 2020) and marmosets (Smith et al., 2009). Because we were testing the effect of IN OXT on this early phase of a female-male introduction, we predicted that IN OXT would reduce the escalation to contact aggression but also increase affiliative behavior as described in the introduction. We found similar levels of lunging and chasing behavior in both OXT and control males, but control males engaged in more wrestling aggression, leading to a significantly higher proportion of control males that escalated their aggression to contact aggression. In this context, OXT may increase the rapid social assessment of and approach towards a potential mate, attenuating high levels of aggression. This change in behavior may decrease time to pair bonding and reduce the chance of injury because males are approaching females with less intense aggression. In the time frame of this test, we did not see a transition to affiliative behavior in either OXT or control males. Similar OXT-driven reductions of aggression in mating contexts have been observed in female Syrian hamsters (Harmon et al., 2002). However, this is the first study reporting anti-aggressive effects of OXT during intersexual interactions in males towards females. This anti-aggressive effect of OXT may have been revealed in California mice specifically because they are a highly aggressive species that also has a prolonged courtship phase prior to mating.

In contrast to opposite-sex social interactions, encounters with unfamiliar individuals of the same sex interactions do not have the same potential for affiliative behavior in a highly monogamous and territorial species. While we predicted that IN OXT would increase escalation to contact aggression in the resident-intruder paradigm, we found that there was no difference in aggression between control and IN OXT treated males. This is consistent with another study that found the same dose of IN OXT used in this study (0.8 IU/kg) did not influence numbers of bites or attack latency in a resident intruder aggression test in California mice (Steinman et al., 2016). It is possible that in a highly territorial and monogamous species there may be selection for a maximum aggressive response to an intruding male. Interestingly, intracerebroventricular injections of vasopressin increased did not increase aggression in a resident-intruder paradigm for male California mice, but a V1a antagonist decreased aggression, further supporting the idea of a maximum level of aggression (Bester-Meredith et al., 2005). Previous studies in less territorial species have found that OXT increases aggression. In house mice, OXTR null mice expressed increased intrasexual aggression (Devries et al., 1997). A study in female rats that manipulated OXT in lateral septum demonstrates that OXT increases and vasopressin decreases aggression towards same-sex intruders (Oliveira et al., 2021). Studies in humans have also shown an association between increased aggression, competition, and OXT (DeWall et al., 2014; Ne’eman et al., 2016; De Dreu, 2012; Fischer-Shofty et al., 2013). However, studies in monogamous marmosets (Cavanaugh et al., 2018), monogamous titi monkeys (Witczak et al., 2018), female and male rats (De Jong et al., 2014; Calcagnoli et al., 2013; Calcagnoli et al., 2015a; Calcagnoli et al., 2015b), house mice primed for aggressive behavior due to social isolation (Tan et al., 2019), and house mice bred for callous traits (Zoratto et al., 2018) found that OXT was associated with reduced intrasexual competition and aggression. Together with our data, these findings suggest that OXT’s effect on intrasexual aggression may depend heavily on the species, brain areas activated by OXT, and social context.

In our last test, we aimed to assess whether IN OXT had similar prosocial effects in fathers as it did in California mice mothers (Guoynes & Marler, 2021). We predicted a positive prosocial effect on both paternal behavior and vocalizations. We found that IN OXT decreased paternal latency to approach their pups but did not influence overall level of paternal care. Studies in Mandarin voles have also shown similar effects of OXT on latency to engage in paternal care (Yuan et al., 2019). Reduced latency to approach pups in IN OXT fathers suggests that IN OXT may increase paternal motivation for pup contact without altering the quality of paternal care. This is supported by studies that show activation of the OXT system can increase dopamine and reinforce rewarding behavior (Borland et al., 2018; Borland et al., 2019; Dolen et al., 2013; Martins et al., 2021). However, it is also possible that the decreased latency to approach pups was driven by dampening anxiety during the challenge test. Several studies have also shown that OXT can reduce anxiety and facilitate prosocial approach (Steinman et al., 2019; Williams et al., 2020; Cohen & Shamay-Tsoory, 2018; Domes et al., 2019). Because we did not observe any overall differences in level of paternal care during the test, the effects of OXT on paternal care may be rapid and more likely to influence paternal responsiveness in California mice versus quality of paternal care seen in marmosets (Saito & Nakamura, 2011; Finkenwirth et al., 2016) and human fathers (Naber et al., 2010; Feldman et al., 2010; Gordon et al., 2017; Li et al., 2017; review by Guoynes & Marler, 2020). We again see species variation in the effect of OXT on paternal care, suggesting that differences across species and brain connectivity may have significant impacts on the how OXT will affect paternal care.

In contrast to the positive association between simple sweeps and maternal care, simple sweeps produced by fathers did not have any relationship with paternal care. This could be due to fathers producing a lower number of calls than mothers during the same testing time frame (mothers produced approximately 1.0 simple sweep/s compared to fathers that produced approximately 0.33 simple sweeps/s) (Guoynes & Marler, 2021). However, it is also possible that fathers are more stressed in the absence of their partner than mothers are and therefore vocalize less. This is supported by findings in several other species that show blunted vocalization in response to heighted stress (Lumley et al., 1999; Chabout et al., 2012; Simola & Granon, 2019; Riaz et al., 2015). Lastly, it is also possible that there are sex differences in the function of simple sweeps in California mice, and that mothers rely more heavily on this call than fathers. Previous research in the lab has shown that while both fathers and mothers show biparental care, there are differences in parental care expression between fathers and mothers.

For example, during a very similar paradigm, mothers showed retrieval behavior, unlike fathers in this test (Guoynes & Marler, 2021), and when both parents are together and given a resident intruder challenge in the presence of their pups, fathers were first to approach pups while mothers did significantly more retrieving behavior (Rieger et al., 2019). This suggests that fathers and mothers may divide parental care duties differently and may, therefore, vocalize and communicate differently.

Overall, the social challenges tested during these experiments show that IN OXT increases prosocial approach behavior in affiliative-prone contexts, but not during the context of direct threat or competition. These results align with the social salience hypothesis of OXT (Kemp & Guastella, 2010; Shamay-Tsoory & Abu-Akel, 2016; Peled-Avron & Shamay-Tsoory, 2018). This hypothesis suggests OXT enhances the processing of social stimuli and that this can either lead to affiliative or aggressive behavior depending on the environment, social stimuli, and internal state of the animal. Across the lifespan in a monogamous, territorial species, it is critical to assess social contexts and balance the costs of aggression and challenges with the benefits of mating opportunities and offspring-rearing. To our knowledge, our study is the first to assess the effect of IN OXT during different life-stage challenges in the same animal. Furthermore, our study was the first to show an effect of OXT dampening aggression during pre-courtship female-male interactions.

## Acknowledgements

We would like to thank NSF grant IOS-1946613 for generously funding this research, undergraduate student support for their work during implementation of the experiment and quantification of behavior, UW Madison animal care staff for their excellent care of the animals, and the Serendipity Scholarship Award for summer funding.

