## Supplemental Figure 1 for "Intranasal oxytocin reduces pre-courtship aggression and increases paternal response in California mice (*Peromyscus californicus*)"

| **Behavior** | **Type of test behavior was measured in** | **Description** |
| --- | --- | --- |
| *Lunging* | Pre-courtship, resident intruder | When the mouse lifts both front paws up and quickly extends them out toward another mouse. |
| *Chasing* | Pre-courtship, resident intruder | When the mouse runs behind the stimulus mouse at a high rate of speed and follows with less than a tail’s length distance. |
| *Wrestling* | Pre-courtship, resident intruder | When the mouse and the stimulus mouse have physical contact accompanied by at least two of the following: putting body weight on top of the other mouse, tail thrashing, biting, audible vocalizations. |
| *Body sniffing* | Pre-courtship, resident intruder | Sniffing that occurs within a whisker’s length of segment of the body other than the anogenital region. |
| *Anogenital sniffing* | Pre-courtship, resident intruder | Sniffing that occurs within a whisker’s length of the anogenital region of another mouse. |
| *Autogrooming* | Pre-courtship, resident intruder, paternal care | Licking, biting, or scratching their own body, at any location. |
| *Rearing* | Pre-courtship, resident intruder, paternal care | Both front paws lift up towards the wall of the cage. Counted as an additional event if front paws touch the ground again and lift back up. |
| *Freezing* | Paternal care | When the mouse does not move at all but has eyes alert/open. Usually ears are up, and they do not make any other movements. Considered a second event once mouse moves for at least 2 seconds and before freezing again. |
| *Latency to approach pups* | Paternal care | Amount of time that it takes the father to first reach his pups after the partitioned mesh door is opened. |
| *Huddling* | Paternal care | Amount of time that the father spends over the pups. At least 50% of the body of pup must be covered by the father to count. |
| *Licking and grooming* | Paternal care | Amount of time that the father spends licking and grooming pups. |
| *Retrieving/carrying* | Paternal care | Amount of time that the father spends with a pup in his mouth while locomoting in the chamber. It is counted regardless of whether locomotion is toward or away from home bedding and cotton ball. |

**S. Figure 1. Ethogram with description of behaviors measured in each test.**
