## Supplemental Table 1 for "Intranasal oxytocin reduces pre-courtship aggression and increases paternal response in California mice (*Peromyscus californicus*)"

| **Behavior** | **Treatment** | **Mean** | **SEM** |
| --- | --- | --- | --- |
| *Lunging* | CTRL  OXT | 1.29  0.45 | ± 1.36  ± 0.37 |
| *Chasing* | CTRL  OXT | 10.76  12.80 | ± 3.73  ± 3.82 |
| *Wrestling* | CTRL  OXT | 11.58  0.77 | ± 6.22  ± 0.50 |
| *Body sniff* | CTRL  OXT | 36.38  58.59 | ± 10.00  ± 10.90 |
| *Anogenital sniff* | CTRL  OXT | 38.33  47.82 | ± 11.22  ± 12.74 |
| *Autogrooming* | CTRL  OXT | 22.08  26.09 | ± 8.55  ± 8.50 |
| *Rearing* | CTRL  OXT | 60.42  73.91 | ± 11.93  ± 18.24 |

**S. Table 1.** All behaviors measured during the pre-courtship aggression test.
