## Supplemental Table 2 for "Intranasal oxytocin reduces pre-courtship aggression and increases paternal response in California mice (*Peromyscus californicus*)"

| **Behavior** | **Treatment** | **Mean** | **SEM** |
| --- | --- | --- | --- |
| *Lunging* | CTRL  OXT | 2.25  1.63 | ± 1.00  ± 0.71 |
| *Chasing* | CTRL  OXT | 13.63  11.28 | ± 5.52  ± 5.86 |
| *Wrestling* | CTRL  OXT | 11.47  10.69 | ± 4.52  ± 4.63 |
| *Body sniff* | CTRL  OXT | 16.11  20.39 | ± 8.31  ± 3.75 |
| *Anogenital sniff* | CTRL  OXT | 29.71  65.87 | ± 9.58  ± 16.73 |
| *Autogrooming* | CTRL  OXT | 2.22  4.24 | ± 0.97  ± 1.60 |
| *Freezing* | CTRL  OXT | 3.02  0.59 | ± 2.84  ± 0.72 |
| *Rearing* | CTRL  OXT | 24.04  26.96 | ± 13.04  ± 15.32 |

**S. Table 2.** All behaviors measured during the resident intruder aggression test.
