## Supplemental Table 3 for "Intranasal oxytocin reduces pre-courtship aggression and increases paternal response in California mice (*Peromyscus californicus*)"

| **Behavior** | **Treatment** | **Mean** | **SEM** |
| --- | --- | --- | --- |
| *Number of pups* | CTRL  OXT | 2.13  2.25 | ± 1.84  ± 1.54 |
| *Licking and grooming* | CTRL  OXT | 34.50  54.88 | ± 15.80  ± 13.91 |
| *Retrieving/carrying* | CTRL  OXT | 0  0 | ± 0  ± 0 |
| *Freezing* | CTRL  OXT | 39.94  8.00 | ± 14.73  ± 4.57 |
| *Rearing* | CTRL  OXT | 26.58  30.38 | ± 10.74  ± 8.79 |
| *Autogrooming* | CTRL  OXT | 3.13  10.19 | ± 1.69  ± 3.42 |

**S. Table 3.** Behaviors measure but not statistically analyzed during the paternal care test.
